# Fast-ripples are emergent properties of neuronal networks

**DOI:** 10.64898/2026.02.15.706005

**Authors:** Laurent Sheybani, Yichen Qiu, Prince Kumar Singh, Umesh Vivekananda, Neil Burgess, Beate Diehl, Andrew McEvoy, Anna Miserocchi, James A. Bisby, Tawfeeq Shekh-Ahmad, Gabriele Lignani, Daniel Bush, Matthew C Walker

## Abstract

Fast-ripples (250-500 Hz) have been proposed as a promising biomarker in epilepsy, but their specificity remains unclear. In particular, it is uncertain whether they reflect chance coincident neural activity or distinctly generated pathological entities. We combined *in silico* simulations, neuronal cultures, the intra-hippocampal kainate rat model of hippocampal epilepsy, and human microwire recordings to investigate whether fast-ripples occur more frequently than expected by the chance aggregation of action potentials. Our simulations showed that chance aggregation can generate fast-ripples and that their incidence changes depending on brain state, an observation that we confirmed in our rodent data. The likelihood of exceeding chance depended on biological system complexity and vigilance state: fast-ripples in neuronal cultures did not surpass chance levels, whereas those in awake – but not sleeping – rodents did. Similarly, the incidence of fast-ripples in awake human recordings was ∼30% greater than expected by chance. As such, our findings suggest that most fast-ripples reflect stochastic network activity rather than distinctly generated pathological entities. This work does not rule out FRs as potential indicators of epileptogenic tissue, but it does challenge prevailing assumptions about their generation and specificity. Their higher prevalence in epileptogenic tissue is likely primarily due to increased excitation and/or neural synchronization, rather than peculiar abnormalities in network behavior.

## Introduction

In epilepsy, disease biomarkers are central to diagnosis and treatment, in particular the decision on where to operate (Duncan et al., 2006; Ewen et al., 2025; Zijlmans et al., 2019). Two main classes of biomarker can be defined. On the one hand, some biomarkers share the same generative mechanisms as the disease itself, making them highly valuable and specific. Other biomarkers are epiphenomena that emerge from activity related to, but not specific to, the disease. While they can also be valuable, great care must be taken not to overinterpret their relevance in diagnostic studies. Importantly, in epilepsy, most biomarkers are epiphenomena. For example, interictal epileptiform discharges and neuroimaging findings provide only indirect spatial information on the epileptogenic zone – i.e., the parenchyma that needs removal to obtain seizure freedom (Krumholz et al., 2015; Mégevand et al., 2014; Sheybani et al., 2025; Zijlmans et al., 2019).

Another electrophysiological biomarker that has gained considerable attention are fast-ripples (FRs). FRs are high-frequency oscillations that are proposed to be specific to brain regions that generate seizures (Dimakopoulos et al., 2024; Jacobs et al., 2010, 2008; Nevalainen et al., 2020), although conflicting results exist in the literature (Dimakopoulos et al., 2024; Klimes et al., 2019; Lisgaras and Scharfman, 2023; Roehri et al., 2018; Sheybani et al., 2019, 2018; Zijlmans et al., 2012, 2009). FRs are detected in the local-field potential (LFP) at high frequency ranges (lower band: 200-250 Hz (Dimakopoulos et al., 2024; Roehri et al., 2024); higher band: 500-550 Hz (Dimakopoulos et al., 2024; Roehri et al., 2024; Sheybani et al., 2018)). While lower frequency (< 200 Hz) LFP fluctuations primarily relate to post-synaptic, i.e., dendritic, potentials (Cohen, 2017; Staba, 2012), this higher frequency activity is dictated by local action potentials (APs) (Logothetis, 2002; Ray et al., 2008). Indeed, previous studies have consistently demonstrated that FRs reflect sequences of APs generated at a specific frequency or inter-spike interval (Dzhala and Staley, 2004; Foffani et al., 2007; Ibarz et al., 2010; Staba, 2012; Ylinen et al., 1995).

Despite decades of research into FRs (Bragin et al., 1999), it remains unclear whether they represent distinct pathological entities unique to epilepsy or arise from coincident neural activity that may be more prevalent in, but is not specific to, epileptic networks. The recent identification of high-frequency oscillations in neurodegenerative diseases supports this conceptual shift (Lisgaras and Scharfman, 2023). Hence, FRs could emerge when there is chance clustering of APs giving rise to an oscillatory pattern. Alternatively, they could occur as pathological entities distinct from other brain processes, characterized by their own specific mechanisms of generation. Conceptually, this distinction is similar to the finite monkey theorem, which states that letting a monkey press randomly on a typewriter will eventually lead to it writing the entire works of Shakespeare. Hence, FRs could either be the product of a succession of chance clusters of APs (i.e., English letters) or distinct entities (i.e., full English words, Fig. 1a).

**Figure 1.**
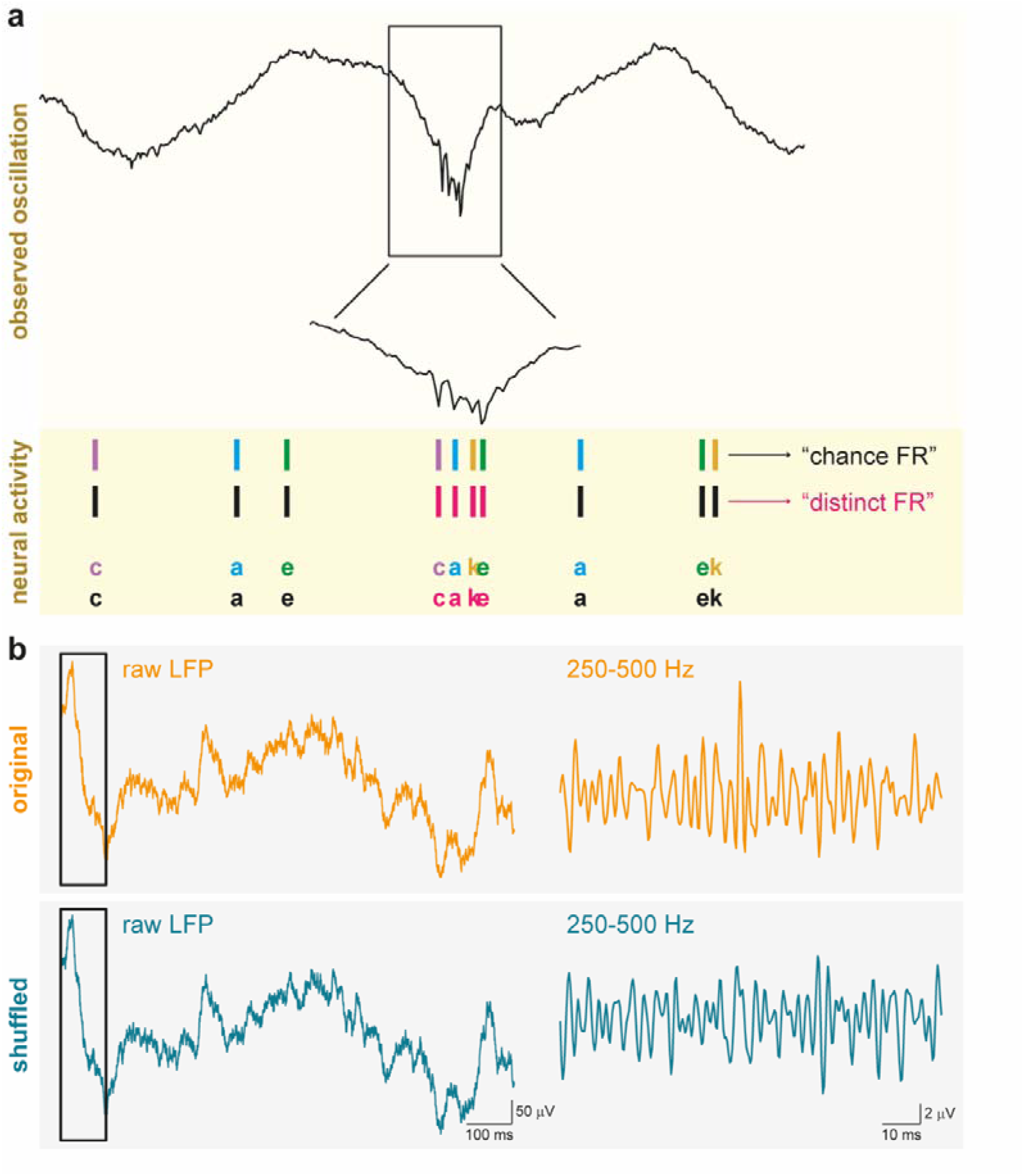
Are FRs “chance activities” or “distinct phenomena”? **(a)** Fast-ripples (FRs) can arise by the chance plesiochronous firing of neurons or can be generated as distinct entities akin to a pathological motif. Similarly, words can emerge either by chance when tapping randomly on a typewriter, or as individual lexical entities. **(b)** To investigate the relative proportion of true and chance FRs, we shuffled EEG signals within the frequency band of interest (250-500 Hz). This disrupts any ‘true’ FRs and therefore allows us to estimate the incidence of chance FRs. The black rectangles on the left are filtered and expanded on the right.

Here, using an *in silico* model, multi-electrode array (MEA) recordings of neuronal cultures, the kainate rat model of temporal lobe epilepsy, and human microelectrode recordings, we asked whether FRs are distinct phenomena or result from the chance clustering of APs. This is central to understand whether the high incidence of FRs in the epileptogenic zone (Dimakopoulos et al., 2024; Jacobs et al., 2008) is simply a byproduct of altered network properties such as firing rate and synchronization, or whether the epileptogenic parenchyma displays intrinsic characteristics that generate FRs independent of these properties. Our results show that the majority of FRs in rats and humans occur due to the chance clustering of APs, that this proportion fluctuates along the sleep-wake cycle, and that the occurrence of emergent FRs is constrained by the overall size of the neural population, neural firing rate and synchronization. These results call for a re-evaluation of the role of FRs as specific biomarkers of epilepsy and support their high susceptibility to changes in neural excitability. This contrasts with the view that they mainly represent specific products of the epileptogenic parenchyma, independent from network properties (Bragin et al., 2000).

## Results

### FRs can occur as emergent phenomena *in silico*

In the finite monkey theorem, it has been calculated that the universe would most likely end before the entire works of Shakespeare were written (Woodcock and Falletta, 2024). Similarly, it could be that the mean firing rate of a neural population and its level of synchronization are such that it very rarely leads to emergent FRs within the duration of an EEG recording or human lifespan. To test this, we first constructed a simulation based on background activity sampled from a human intracranial microelectrode recording, and inserted APs arising from incremental numbers of neurons (population of 1-2000 neurons, 50 steps), with progressively increasing mean firing rates (1-20 Hz, 50 steps) and level of synchronization (50 steps exponentially related to the Fano factor, a measure of the dispersion of neural discharges, Fig. 2a and Supplementary Fig. 1). The model confirmed that FRs can emerge by chance, i.e., as a result of randomly inserting APs (Fig. 2b), and displayed the expected spectral signature of FRs (Fig. 2c-d). Similar approaches have been described previously (Zanos et al., 2011), building on the observation that high frequency activity in the LFP reflects neuronal discharges (Fedele et al., 2020; Manning et al., 2009; Quiroga et al., 2004; Ray et al., 2008) and that neuronal discharges are the building blocks of FRs (Foffani et al., 2007; Ibarz et al., 2010; Staba, 2012).

**Figure 2.**
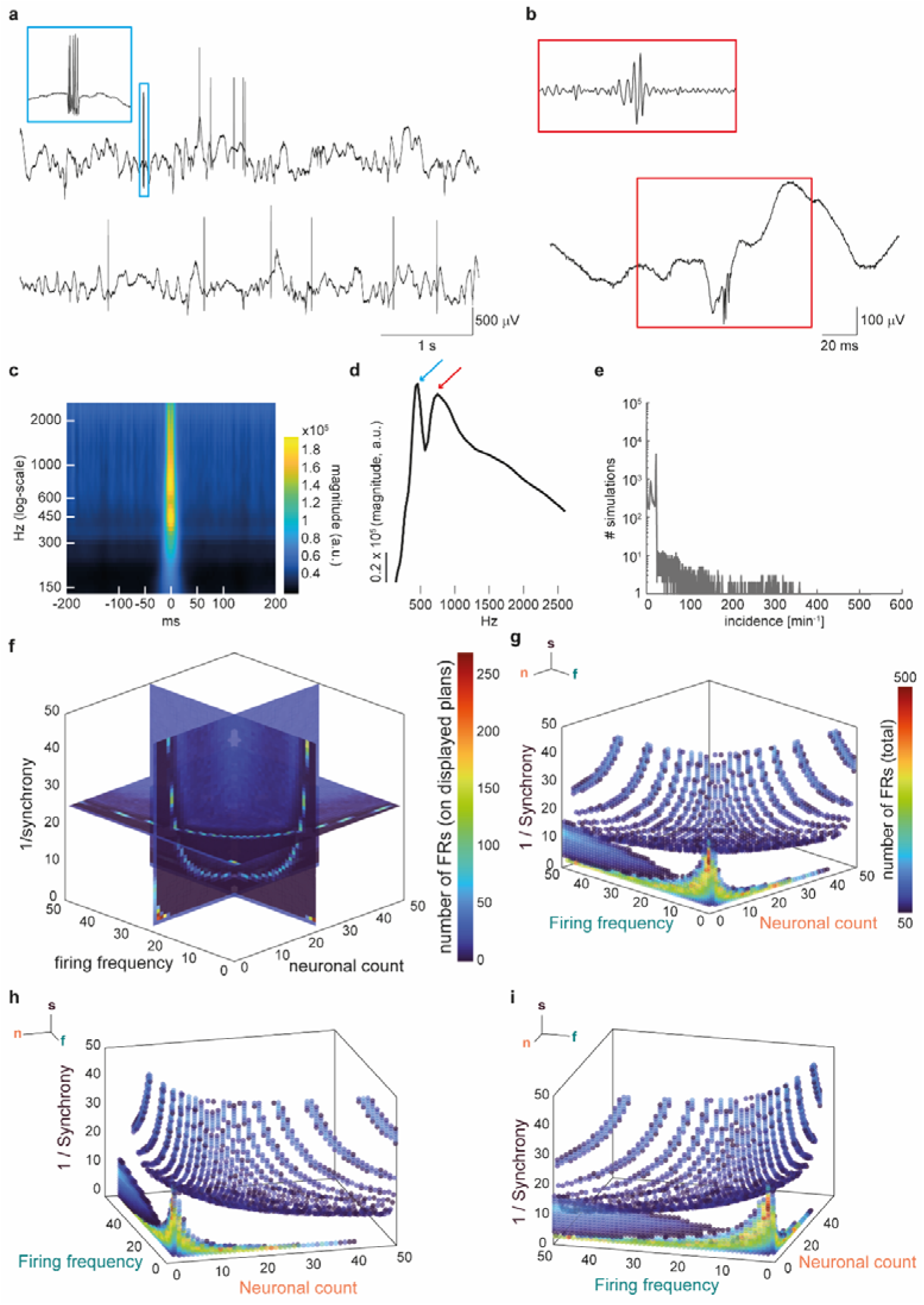
Fast-ripples (FRs) can arise from stochastic neural firing. **(a)** Example of 5-s activity from 2 signals with inserted action potentials (APs). Across their entire length (480-s), both signals contain a similar number of APs but the upper trace displays more synchrony than the lower trace, where firing is even. The blue squares highlight a burst of highly synchronized APs. **(b)** Identification of FRs in simulated signals. **(c)** Time-frequency representation of detected FRs in simulated signals. **(d)** Spectrogram of detected FRs in simulated signals. The left peak (blue arrow) reflects FRs and the right peak (red arrow) reflects individual APs. **(e)** Number of simulations (y-axis) per FRs rate (x-axis). **(f)** Number of detected FRs across all simulations (50x50x50=125,000) with varying numbers of neurons, mean firing frequency, and synchronization. FRs do not occur randomly; a pattern can be identified, indicating that these three parameters control the expression of FRs. **(g-i)** Solution points for > 50 FRs to better highlight the pattern of expression. n=neuronal count, f=firing rate, s=synchronization.

### Emergent FRs index neuronal population, firing rate and synchronization

Different simulations, i.e., with different neuronal count, firing frequency and synchronization parameters, could lead to the same incidence of FRs (Fig. 2e). Hence, we next tested whether changes in the incidence of FRs was random across simulations or whether it could be predicted by changes in neuronal count, firing frequency and/or synchronization. An absence of stochasticity would imply that FR incidence is dictated by one, or a combination of, these parameters.

We found that across the space of 125,000 (i.e., 50*50*50) parameter values, we observed a structured pattern in the number of detected FRs (Fig. 2f-i). We then sought to determine if this pattern was constrained by neuronal count, firing rate and synchronization. To do so, we ran a gradient boosted tree (GBT) to predict the number of detected FRs based on these three features. The GBT predicted a distribution of FR densities which was similar to the true density of FRs (Fig. 3a-b) across solution points (*r^2^*=0.73, *p*<0.0001, Fig. 3c). When shuffling the three parameters (neuronal count, firing rate, synchronization), we found that the explained variance (R^2^) was significantly lower than the variance explained by the original GBT model (mean ± standard deviation [SD] at chance level: 0.02 ± 0.001, *p*=0.002, Fig. 3d). Furthermore, using a cross-validation approach with 80% of the data as training set and the remaining 20% as the test set, we obtained a significantly higher explained variance than when outputs were shuffled across the 125,000 solution points (paired t-test, *p*<0.0001, Fig. 3e), confirming that neuronal count, firing rate and synchronization are significant predictors of the incidence of FRs. Importantly, this analysis excludes the possibility that detected FRs were only those already present in the background microelectrode signal, which was fixed across all simulations. Hence, not only can FRs emerge due to coincident neural firing, but their incidence indexes the size of the neuronal population, its mean firing rate and level of synchrony.

**Figure 3.**
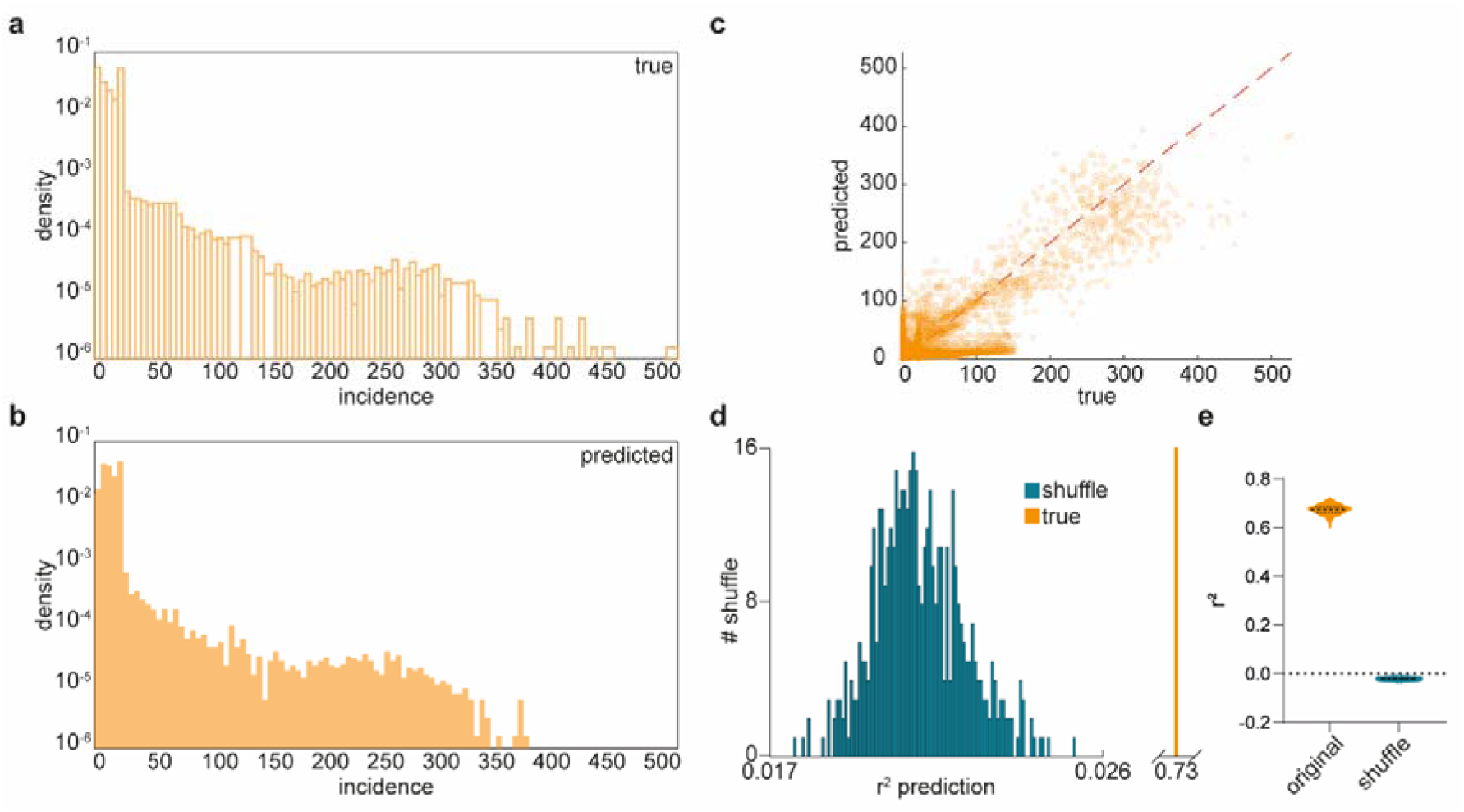
Neural count, firing frequency and synchronization constrain the emergence of FRs. **(a)** Density of FRs across the 125,000 solution points of the simulation. **(b)** Density of FRs across the 125,000 solution points predicted by a gradient boosted tree (GBT). The distribution is very similar to (a). **(c)** All solution points predicted by the GBT (y-axis) along corresponding solution points of the original simulation (x-axis). The correlation is significant (*r^2^*=0.73, *p*<0.0001). **(d)** *Blue* Performance of the GBT to predict FR number, should FRs be randomly distributed across values of neural count, firing frequency and synchronization (500 permutations). *Orange* Performance of the GBT to predict FR number in actual simulations. **(e)** Performance of the GBT classifier using cross-validation (training: 80% of data; test: 20% remaining) using original (orange) and shuffled (blue) data. The difference is significant (paired t-test, *p*<0.0001).

### FR rate is at chance level in neuronal cultures

The model described above predicts that FRs can occur as the result of chance firing. We hypothesized that in a simple biological system such as a neuronal culture, FRs would not be more frequent than this expected “chance rate”. To test this, we recorded the activity of neural cultures using MEAs under a baseline condition and after administration of the GABA_A_ receptor antagonist picrotoxin, a pro-seizure compound that increases network excitability and the generation of epileptiform activity (Supplementary Fig. 2). We hypothesized that if FRs arise purely from chance firing, then temporally shuffling these recordings while conserving their spectral properties (Supplementary Fig. 3) would disrupt any oscillatory structure, leaving only FRs that occur due to chance. Any additional FRs in the original data, compared to the number of FRs in the shuffled EEG, should thus be assumed to be individual entities. We identified FRs in original signals (Fig. 4a), but they were not significantly more frequent than in the shuffled signals (main effect of shuffling: F(1,15)=0.99, *p*=0.33, Fig. 4b). In addition, we did not identify a main effect of picrotoxin (F(1,15)=1.53, *p*=0.24) or shuffling * picrotoxin interaction (F(1,15)=1.11, *p*=0.31). Hence, neural networks with limited complexity (Kim et al., 2020; Saglam-Metiner et al., 2024; Sanchez-Vives and McCormick, 2000; Timofeev and Chauvette, 2017) fail to generate FRs beyond that expected from the chance coincidence of APs, even after increasing network excitability.

**Figure 4.**
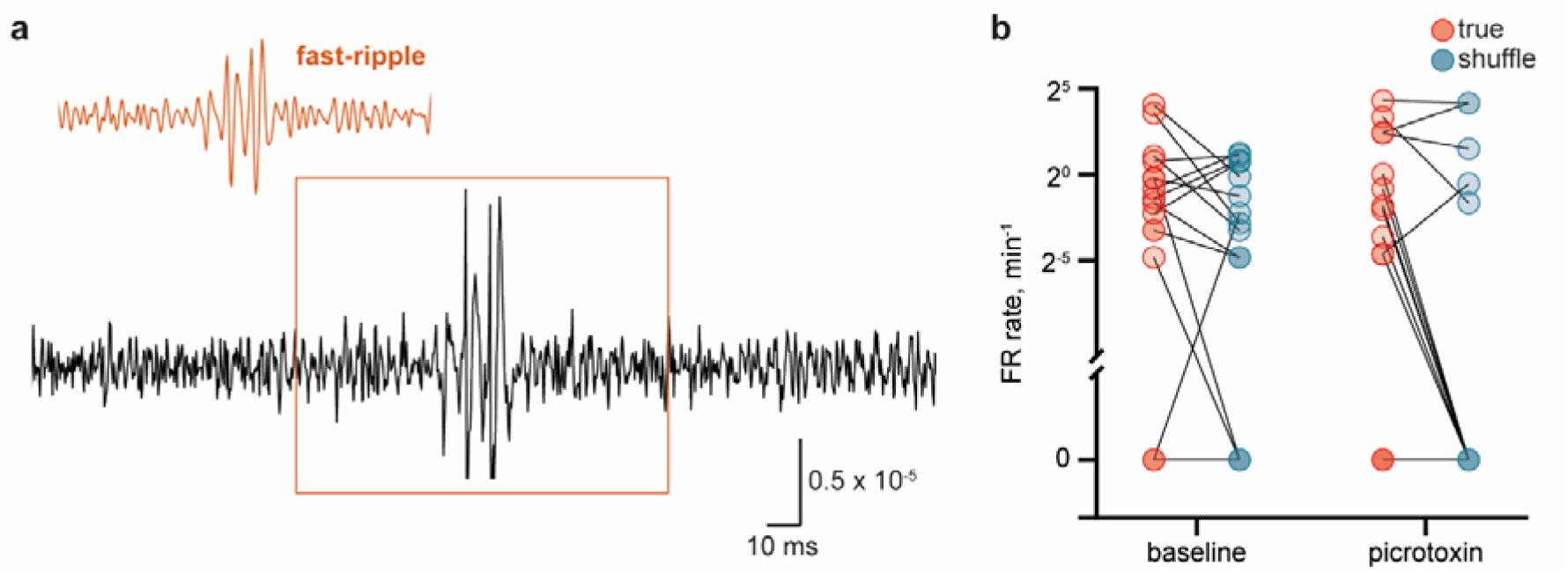
FRs in neural culture. **(a)** Example of a detected FR in an MEA recording of neural culture. **(b)** In the MEA recordings, there are no more FRs than expected by chance (‘shuffle’) during baseline or in a pro-epileptic state following application of picrotoxin.

### FR rate is predicted by neural firing rate and synchronization

Next, we used a rodent model of temporal lobe epilepsy to study the emergence of FRs *in vivo*. Across week-long recordings, and in line with the known variability of FRs across time (Gliske et al., 2018), we observed a diurnal variation in FR rate (Fig. 5a-c, orange trace). By comparing FR rate to delta power, we could infer how FRs evolve across different vigilance states, assuming that high and low delta activity reflect sleep and wake periods respectively (Osorio-Forero et al., 2024). FRs were more frequent during periods of presumed wakefulness: indeed, FR incidence was strongly modulated by delta activity (Supplementary Fig. 4a) and peaked at delta power troughs (Supplementary Fig. 4b). Importantly, these findings were robust to the specific method used to detect FRs (Supplementary Fig. 4c) (Padmasola et al., 2024; Sheybani et al., 2019, 2018). Also our use of delta power to identify periods of presumed wakefulness and sleep was highly (negatively) correlated with an alternative method of using beta to delta ratio across time (another marker of increased vigilance; (Fraigne et al., 2023), see Supplementary Fig. 4d). Furthermore, we confirmed that the higher the incidence of FRs during wakefulness was not due to movement artifacts (Bénar et al., 2010). Indeed, FRs were visually concordant with those described in the literature (Supplementary Fig. 5) and not associated with surges in broadband high frequency activity that might reflect muscle artifacts (Supplementary Fig. 6).

**Figure 5.**
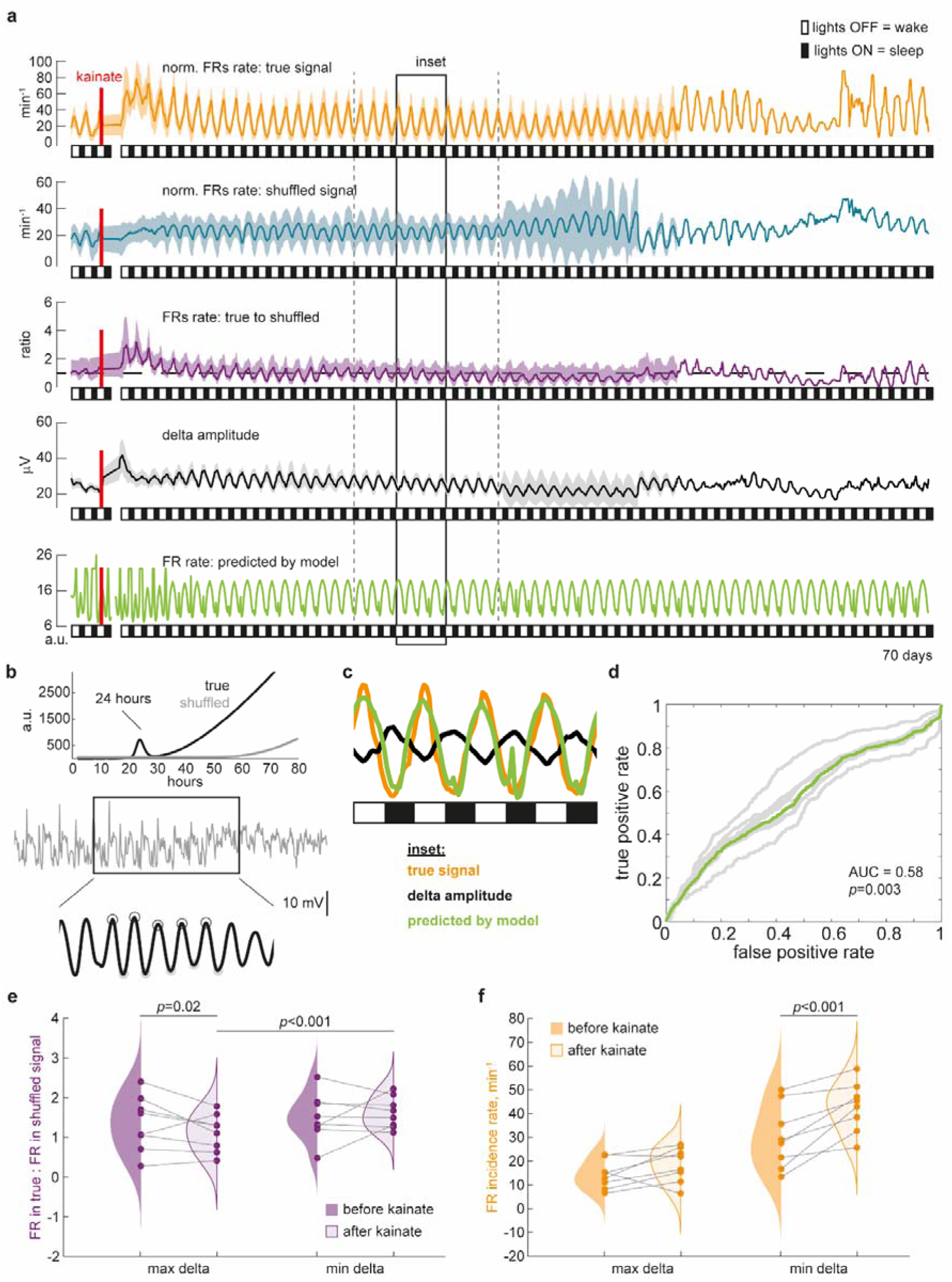
The proportion of FRs as distinct entities varies across the sleep-wake cycle. **(a)** Incidence of FRs in the original signal (orange trace), shuffled signal (blue), the ratio of FR incidence in the original to shuffled signals (purple), delta amplitude (black) and predicted incidence of FRs across time. Here and after, these metrics are sampled at a rate of h^-1^. **(b)** Periodogram of delta amplitude across days. A clear peak at 24 hours is seen in original (black) but not shuffled (grey) data. **(c)** Fluctuation of FR incidence across days in the data (orange) and as predicted by the model (green), overlaid on fluctuations in delta amplitude. **(d)** AUC of the model to predict high (above median) and low (below median of individual rats’ FR incidence rate) values of FR incidence rate. The AUC is significantly above chance level, i.e., 0.5 (mean±SD: 0.58±0.05, *p*=0.003, one-sampled t-test). Each grey line is one animal, the green line is the average across animals. **(e)** Two-way ANOVA comparing the ratio of FR incidence rate in original to shuffled signals, before and after kainate injection, sampled at delta peaks (max delta) and troughs (min delta). There is a significant kainate * delta interaction (F(1,7)=17.1, *p*=0.004) and post-hoc tests show that the ratio is different before and after kainate injection only at delta peaks (mean difference, 95%: 0.24, [0.10-0.38], *p*=0.02). **(f)** Two-way ANOVA comparing the incidence rate of FRs before and after kainate injection, sampled at delta peaks and troughs. We observed a significant kainate * delta interaction (F(1,7)=14.7, *p*=0.006). The incidence rate after kainate injection increased only when sampled at delta troughs (12.6 min^-1^, [9.0-16.3], *p*<0.001) and not peaks (4.2 min^-1^, [0.5-7.9], *p*=0.12).

Given our simulation results, we hypothesised that this diurnal variance may arise from fluctuations in firing rates and synchronization, which are known to decrease and to increase during sleep, respectively (Frauscher et al., 2016; Miyawaki et al., 2019; Staba et al., 2002; Watson et al., 2016). To test this, we first established the peaks of FR rate variability, which, as expected, were identified at 24-hour intervals (Supplementary Fig. 7). Based on this period, we then randomly sampled different simulations to generate synthetic results at hourly intervals over 70 days, with the only constraint being that firing rate decreases and synchronization increases during sleep (Frauscher et al., 2016; Miyawaki et al., 2019; Staba et al., 2002; Watson et al., 2016). Sleep was defined as a 12-hour period surrounding the maxima of delta activity, which also peaked at 24-hour intervals (Fig. 5b). Using only these two constraints, the model exhibited a daily fluctuation of FR rate, which was higher during wake than sleep (Fig. 5a and 5c, green trace). The actual rate of FRs was correlated significantly more with the predicted rate (median *r^2^* coefficient, IQ range: 0.01, [0.004-0.04]) than with a shuffled version of the predicted rate (0.0009, [0.0008-0.001], *p*=0.016, Wilcoxon test) and the ability of the model to predict periods of high and low incidence rate of FRs was significant (mean AUC ± SD: 0.58, ± 0.05, *p*=0.003, one-sample t-test, Fig. 5d). Furthermore, the periodogram of FR incidence in simulated EEGs also displayed a clear peak at 24 hours (Supplementary Fig. 8). Hence, changes in neural firing rate and synchronization across the sleep-wake cycle can explain the increased incidence of FRs observed in rodents during wakefulness *in vivo*.

### The relative proportion of distinct and chance FRs depends on brain state

That neural firing rate and synchronization modulate chance FR incidence does not exclude the possibility that distinct FRs may still occur. To determine the proportion of distinct FRs in our chronic *in vivo* recordings, we again computed the incidence of FRs in original and temporally shuffled signals. Similar to the rate of FRs in the original signal, we observed a daily fluctuation of FRs in the shuffled signal (Fig. 5a, blue trace) and in the ratio of FR rate in the original to shuffled signal (Fig. 5a, purple trace). We then tested if this ratio varies depending on brain state, i.e., sleep vs wake. As already shown, and similar to FR rate, delta activity fluctuated with a period of 24 hours, reflecting sleep periods (Fig. 5b, *top*). We filtered delta activity at the corresponding frequency (Fig. 5b, *middle* and *bottom*) and identified peaks and troughs of this filtered signal (Fig. 5b, open and grey disks). We then compared the ratio of FR rates (original : shuffled) at these timepoints using a 2-way ANOVA (condition kainate: before vs after; condition delta: at peaks vs at troughs). We found a significant kainate * delta interaction (F(1,7)=17.1, *p*=0.004, Fig. 5e). Post-hoc testing showed that the ratio of original : shuffle FR rates was higher at delta troughs than peaks after kainate injection (0.51, [0.37-0.65], adjusted *p*<0.001), but not before (mean difference, 95% CI: 0.16, [0.02-0.30], adjusted *p*=0.12). Importantly, the ratio decreased at delta peaks after kainate injection (0.24, [0.10-0.38], adjusted *p*=0.02, Fig. 5e) but not at delta troughs (-0.11, [-0.25 to 0.03], adjusted *p*=0.41, Fig. 5e). Furthermore, it shows that the probability of observing FRs that arise purely from the chance clustering of APs varies with brain state.

Last, we found that before kainate injection, the FR rate ratio was not greater than 1, whether it was computed at delta power troughs (1.5, [1-2], adjusted *p*=0.09) or peaks (1.3, [0.8-1.9], adjusted *p*=0.41). Hence, in non-epileptic animals, FRs are no more frequent than expected by chance. In contrast, after kainate injection, we observed a FR rate ratio that was significantly greater than one at delta troughs (mean, 95% CI: 1.6, [1.3-2], adjusted *p*=0.006, one-sampled t-test) but not peaks (1.1, [0.7-1.5], adjusted *p*>0.99, one-sampled t-test). This indicates that, in contrast to non-epileptic animals, epileptic animals exhibit more FRs than would be expected by chance, but only during wakefulness. Note that this ratio was significantly different between delta troughs and peaks after kainate (*p*=0.0003, paired t-test). Hence, the specificity of FRs to epilepsy is shaped by vigilance states.

### Diurnal variation of FR rate as a confound in their interpretation

In practice, clinicians calculate the absolute incidence of FRs, and not the ratio of original to shuffled FRs, as examined above. Hence, we next asked whether the absolute rate of FRs is also impacted by the sleep-wake cycle. Indeed, the periodicity of FR rate could contribute to discrepant interpretations regarding the impact of kainate injection. Visually, while the absolute rate of FRs at troughs of FR incidence seems relatively stable after kainate injection, the FR rate at peaks of FR incidence tends to increase (Fig. 5a, orange). To assess this statistically, we computed a 2-way ANOVA, comparing the absolute incidence rate of FRs before and after kainate injection at the peaks and troughs of delta amplitude identified previously (Fig. 5b), and found a significant kainate * delta interaction (F(1,7)=14.7, *p*=0.006, Fig. 5f). Post-hoc testing showed that FR rate was higher after than before kainate injection at delta power troughs (mean difference, 95% CI: 12.6 min^-1^, [9.0 to 16.3], adjusted *p*<0.001), but not at delta peaks (4.2 min^-1^, [0.5 to 7.9], adjusted *p*=0.12). This confirms that changes in FR incidence after induction of epilepsy varies across the sleep-wake cycle. Specifically, the absolute rate of FRs increases after kainate injection during wakefulness (Fig. 5f). However, the absence of a change in the ratio of original : shuffled FRs during wakefulness, after kainate injection shown above (Fig. 5e), suggests that this arises from changes in general network excitability, rather than the emergence of FRs as distinct entities.

These findings demonstrate the challenge of identifying distinct FRs within a composite population of distinct and stochastic events. One parameter that could help disentangle these events is their duration. Indeed, one might expect stochastic events to be more likely to be short-lived, since the probability of consecutive APs continuing to co-occur across neurons decreases over time. Hence, we next compared the distribution of FR durations between original and shuffled rodent data and found that FRs in shuffled data are shorter than those in original data (Supplementary Fig. 9). This makes duration a key feature that could help identify distinctly generated FRs.

### The majority of FRs also occur by chance during wakefulness in humans

Finally, we asked whether FRs in the human brain were more frequent than expected by chance. To do so, we examined intracranial microelectrode recordings from 10 patients who underwent presurgical epilepsy evaluation. Microwires were implanted in the mesial temporal lobe in all subjects and recordings were obtained during wakefulness. We proceeded again by shuffling the original data and then identifying FRs in the original and shuffled signals (Fig. 6a).

**Figure 6.**
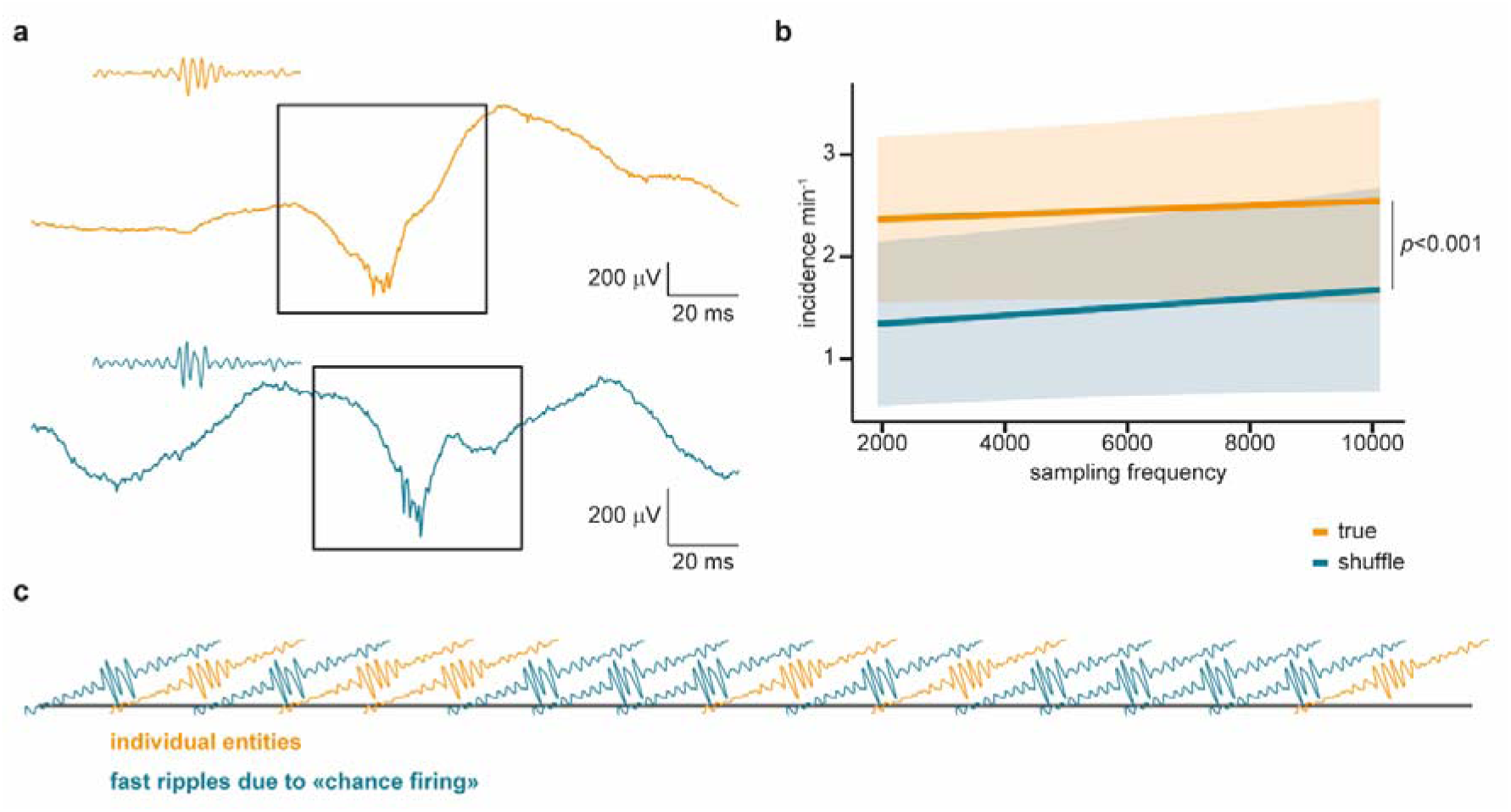
FRs as distinct entities in humans are uncommon. **(a)** Example of a FR detected in the original signal (orange) and shuffled signal (blue). **(b)** Incidence of FRs per minute in the original and shuffled signals, across sampling frequencies (3,000; 4,000; 5,000; 6,000; 8,000; 10,000; 15,000 Hz) to assess any possible effect of sampling frequency. We observed only a main effect of signal type (original vs shuffle, F(1, 770.4)=20.62, *p*<0.001) but no interaction (F(1,770.4)=0.05, *p*=0.82) or main effect of sampling frequency (F(1,13.2)=0.47, *p*=0.51). **(c)** Illustration of the proportion of FRs in the original signal and in the corresponding shuffled signal. This ratio is of 3:5. Hence, distinct FRs account for only 37.5% of all detected FRs.

We found a main effect of signal type (original vs shuffled: F(1,770.4)=20.62, *p*<0.001, Fig. 6b), confirming that FRs are more frequent in the original signal than expected by chance. Specifically, the original recordings displayed ∼1.6 times more FRs than shuffled recordings, such that about 3 FRs can be considered distinct pathological entities for each 5 FRs occurring by chance (8/5 = 1.6), making distinct FRs only 37.5% of all those detected (Fig. 6c). Interestingly, the proportion of distinct to chance FRs in these awake subjects is similar to that seen in awake rats (Fig. 5e). Last, less sophisticated methods that did not restrict the shuffling to the FR frequency band (see Methods) did not display a significantly different rate of FRs in original than in shuffled data (Supplementary Fig. 10). This supports that the main shuffling procedure was not underestimating the proportion of FRs in the original data.

## Discussion

Here, we report that distinct FRs, i.e., those that do not emerge as the chance clustering of APs, account for around a third (37.5%) of all detected FRs during wakefulness in humans. Furthermore, both the incidence rate and relative proportion of distinct FRs vary with brain state (Ewell et al., 2019; Sakuraba et al., 2016), making this electrophysiological entity more complicated to interpret than previously thought (Menendez de la Prida et al., 2015; Roehri et al., 2018). It has been argued that the very high frequency that can be obtained during FRs are due to out-of-phase firing of excitatory neurons (Foffani et al., 2007; Ibarz et al., 2010), which is also consistent with our concept of stochastic firing. The conceptual difference is the degree to which there is any underlying organization of this firing. We argue that in the majority of cases there is no organization, although a substantial minority cannot be explained on a stochastic basis.

In our rodent data, we were initially surprised to find a higher rate of FRs during wakefulness, which contrasts with previous reports in humans (Bagshaw et al., 2009; Staba et al., 2004). However, previous studies only indicate that physiological vs pathological FRs are more easily distinguished during NREM sleep (von Ellenrieder et al., 2016) and that their incidence varies during sleep (Von Ellenrieder et al., 2017), but in hours-long recordings, no differences in incidence have been reported in the mesial temporal lobe (Dümpelmann et al., 2015). Furthermore, the variability of FRs across time (Gliske et al., 2018) indicates that longer nocturnal recordings in humans are necessary. It also suggests that changes in excitability across time could explain this change in FR incidence. Last, but not least, another report did not find a state-dependent expression of FRs in the kainate rat model of temporal lobe epilepsy (Ewell et al., 2019), thus indicating that the variability of FRs across sleep and wake is still an open question, at least in rodents. Hence, the main conclusion on the effect of sleep-wake transitions is that these transitions impact the likelihood of stochastic events, more than dictating the direction (increases vs decreases) of change. It also highlights that the specificity of FRs to epileptogenic parenchyma could vary across the sleep-wake cycle, which would be crucial in epileptology (Dimakopoulos et al., 2024; Roehri et al., 2018; Sheybani et al., 2019, 2018; Zijlmans et al., 2012, 2009). Hence, FRs reflect and are highly susceptible to changes in network excitability. This is consistent with recent findings of HFOs in neurodegenerative diseases (Lisgaras and Scharfman, 2023; Shandilya et al., 2024), where hyperexcitability could potentially be a driver of cognitive decline (Vossel et al., 2016). Similarly, a recent study on ripples (70-180 Hz) reported that these events can largely be attributed to changes in background noise levels (Van Schalkwijk and Helfrich, 2026).

How do we reconcile our findings with reports of specific mechanisms underlying the generation of FRs? Indeed, several candidate mechanisms have been identified, including a necessary role of gap junctions (Curot et al., 2023; Demont Guignard et al., 2012; Draguhn et al., 1998; Dzhala and Staley, 2004; Fabo et al., 2023; Foffani et al., 2007; Köhling and Staley, 2011; Schlingloff et al., 2025; Schmitz et al., 2001; Sheybani et al., 2019; Traub et al., 2001; Valero et al., 2017). However, most studies report mechanisms that are likely or even proven to impact network properties, such as firing rate, synchronization or even neuronal count, which we have shown to contribute to the chance occurrence of FRs. For example, lower ephaptic transmission contributes to decreased spike-timing reliability during sharp-waves ripples and has been suggested to lead to pathological FRs in rats (Foffani et al., 2007). However, this could be driven by decreased neuronal count (Foffani et al., 2007). Even electrical coupling via gap junctions could modulate FR incidence through its impact on synchronization (Bennett and Zukin, 2004; Hürkey et al., 2023). Hence, previous literature may have focused mainly on mechanisms that affect network properties, but not directly FRs.

The highly stereotypical morphology (Sheybani et al., 2018) of FRs, especially in the seizure-onset zone (Liu et al., 2018), is not inconsistent with our findings: our results do not exclude the possibility that some FRs are not consecutive to the chance occurrence of plesiochronous APs. In fact, they support the conclusion that a subset of FRs are distinct entities. The reason why FRs have been relatively successful in the delineation of the seizure-onset zone (Dimakopoulos et al., 2024; Jacobs et al., 2010, 2008; Klimes et al., 2019; Nevalainen et al., 2020; Roehri et al., 2018; Sheybani et al., 2019, 2018; Zweiphenning et al., 2022) could thus reflect the fact that changes in network properties that promote the emergence of FRs are also associated with epileptogenicity; but this does not mean that FRs have their own specific generative mechanisms. This is a crucial difference in our quest to understand the mechanisms underlying FRs, and their coupling with epileptogenic zones.

It is important to note that the results presented here could depend crucially on the specific temporal shuffling approach that we employed, and so several different methods were used to address this concern. Importantly, the algorithm shuffles only the high-frequency component of the signal, bypassing lower frequency activity that could contribute to higher frequency artifacts (although the FR detection algorithm used here should also control for this (Roehri et al., 2018)). Shuffled data exhibits comparable spectral composition to the original signals. Furthermore, our results are supported by six other shuffling procedures, which failed to identify an increased incidence of FRs in physiological data. Hence, we argue that FRs in biological signals are only slightly more frequent than chance, and this can be captured only by the most sophisticated shuffling methods.

Importantly, we do not question the potential of FRs to delineate the seizure-onset zone. Instead, our results suggest that the observed increase in FRs within the epileptogenic zone is an emergent phenomenon – arising due to changes in secondary network properties such as excitability and synchronization, not as a direct result of some pathology that is specific to epilepsy. Depolarizations that promote burst firing should also increase the likelihood of FRs. Paroxysmal depolarization shifts, which are the cellular events underlying epileptiform abnormalities would therefore be expected to co-occur with FRs. Accordingly, epileptiform abnormalities should be associated with increased gamma/high-gamma (> 80 Hz) activity at their source as has been observed in epileptogenic parenchyma (Ren et al., 2015; Sheybani et al., 2021; Weiss et al., 2015). In addition, we show that long-durations FRs are more likely to be distinct oscillations than stochastic events; and so FR duration is a key parameter that should be considered in future studies. Finally, it is important to stress that our study does not suggest that FRs are artefactual. They are distinct oscillations, generated by clusters of APs. Suggesting that FRs are artifacts of the signal is different from stating that FRs appear as a result of the chance clustering of APs within an appropriate time window. The latter is correct and contributes to a significant proportion of FRs. This must be now taken into consideration when studying mechanisms of generation of FRs and seizures, and the role of FRs as disease biomarkers.

Clinically, our findings do not argue for or against the use of FRs as epilepsy biomarkers, as we did not assess their relationship with the epileptogenic zone. However, they do indicate that FRs are not highly specific to epileptic tissue. We would, however, recommend selecting FRs with particularly long duration, which may offer greater specificity and potentially better reliability as a biomarker. The clinical value of FRs as distinct entities or as emergent oscillations remains to be clarified.

## Methods

### Mouse primary neuronal culture recording with multielectrode array (MEA)

Embryonic cortical neuronal cultures were prepared from E18-19 wild type C57BL/6J embryos, plated at density of 50k/well at DIV0 onto recording areas pre-coated with PLL (P2636-25MG) and laminin (L2020-1MG) onto Axion CytoView 24-well plate (M384-tMEA-24W, Axion systems). The MEA system was plated at very high density and this could have led to a ceiling; however, as shown in the simulations, relationship between neural density and FRs rate is neither linear nor monotonic. Each well contains 16 electrodes, covering estimated area 1.1mm x 1.1mm. Activity from the same well shares network properties and culture environment, but this is not fundamentally different from sampling > 1 brain region spatially close to each other. Neurons were maintained at 37 degrees, 5% CO2 with appropriate humidity, with 25% media change every week. Electrophysiological recordings were taken with Mastro Edge system (version 3.7.1.16) where the neurons were maintained at 37 degrees and 5% CO2, for 10-20 minutes per session. Recordings were processed at 12.5 kHz sampling frequency, with low and high pass filter set at 3 kHz and 200 Hz, respectively (Axion Navigator). When PTX (picrotoxin) treatment was applied, 50 mM picrotoxin was dissolved in DMSO, where 50 μM was used for triggering neuronal network activities at DIV20. Baseline data were recorded prior to picrotoxin treatment, and post-picrotoxin treatment recordings were taken 24 hrs afterwards. The analysis of spiking, busting and network burst were carried out with Axion navigator (Axion systems). Raw spiking data was exported with Neural Metric Tool (Axion systems) into csv. format for subsequent analysis. All experiments were carried in compliance with Home Office Project Licence (PP3944290).

### Rat recordings and preprocessing

All experimental protocols were conducted in compliance with the Association for Assessment and Accreditation of Laboratory Animal Care (AAALAC) International standards and were approved by the Institutional Animal Care and Use Committee (IACUC) of the Hebrew University of Jerusalem. This study utilized male Sprague– Dawley rats (150–175 gm; the Hebrew University strain) obtained from the Harlan Laboratories Israel Ltd. (Jerusalem, Israel). All animals were housed under controlled conditions (23 ± 1 °C, 50–60% humidity, 12 h light/dark cycle) with free access to food and water. A minimum acclimatization period of one week was allowed before any experimental procedure to ensure adaptation to the housing environment.

#### Surgical procedure: ECoG transmitter implantation

All animals underwent for implantation of high frequency electrocorticography transmitters (ECoG) under general inhalation anesthesia (induction at 3% isoflurane, maintenance at 1.8–2.3%; Terrell™, USP, Piramal Critical Care). Animals were positioned in a stereotaxic frame (Kopf Instruments, CA, USA) for stereotaxic surgeries. Presurgical analgesia was administered subcutaneously including buprenorphine (0.2 mg/kg; Rich Pharma) and meloxicam (1 mg/kg; Chanelle Pharma). Animals were implanted subcutaneously with a high-frequency single-channel ECoG transmitter (model A3049T5, sampling rate 2048 Hz; Open-Source Instruments, USA) featured with two subdural intracranial electrodes. The recording electrode was placed epidurally above the right dorsal hippocampus (AP −2.8 mm, ML +1.5 mm), while the reference electrode was positioned contralaterally (AP −6.0 to −7.0 mm, ML +3.5 mm). Epidural recordings are highly effective in capturing fast-ripples generated in the hippocampus (Sheybani et al., 2018). The electrodes were secured to the skull with screws and tissue adhesive to ensure stable signal acquisition. Following implantation, the cranial incision was sealed with dental cement. Postoperative care including warmed Ringer’s solution for rehydration and amoxicillin (100 mg/kg; Betamox LA) to prevent post-surgical infection were administered subcutaneously. Animals were monitored until full recovery and returned to their home cages. One to two weeks recovery period was observed before subsequent experiments.

#### Induction of status epilepticus

All rats implanted with ECoG transmitters were subjected to status epilepticus (SE) induction using a single intraperitoneal injection of kainic acid (KA; 10 mg/kg; 5 mg/mL in 0.9% saline; Hello Bio, UK). Following injection, animals were continuously monitored for seizure activity via a wireless ECoG telemetry system coupled with CCTV cameras, enabling real-time acquisition of seizure activity during SE development. SE was behaviorally defined by the experimenter as the onset of sustained class V seizures, according to a modified Racine scale (Racine, 1972), characterized by rearing, forelimb clonus, and loss of posture. Two hours after SE onset, diazepam (10 mg/kg, *i.p.*; Assival®, Teva Pharmaceuticals, Israel) was administered to terminate seizures and minimize mortality. Only animals that exhibited continuous convulsive activity throughout the 2-hour post-SE induction window were included in the study.

#### ECoG data acquisition and Analysis

All animals with subdural ECoG transmitters were housed individually under continuous 24/7 wireless video-ECoG (vECoG) monitoring. ECoG signals were wirelessly transmitted and recorded using Neuroarchiver software (Open-Source Instruments Inc.), enabling high-frequency, high-resolution data acquisition with real-time processing capabilities.

Recorded ECoG traces were segmented into 4-second epochs and analyzed across six quantitative parameters: total power, intermittency, coastline, coherence, asymmetry, and 0.2–640 Hz band power. Each feature was normalized to a 0–1 scale and compared against a user-generated seizure reference library containing validated seizure episodes derived from at least three different animals. Seizure identification was performed using an automated detection algorithm, which classified epochs as seizure-like events if their Euclidean distance from the reference library was <0.2, following established protocols [Open-Source Instruments, Event Detection; (https://www.opensourceinstruments.com/Electronics/A3018/Event_Detection.html).

All automatically detected seizures were subsequently reviewed and confirmed by the blinded investigator. In addition, a subset of detected events was validated using time-synchronized video recordings from cage-mounted CCTV cameras (Microseven). These videos were independently assessed by a second blinded researcher to verify concordance between electrographic seizure activity and behavioral manifestations, thereby ensuring reliable event classification. Among the 10 rats, 8 developed seizures and only epileptic rats were included in analyses. Prior approval was granted by local Ethics Committee (Institutional Animal Care and Use Committee, The Hebrew University of Jerusalem; Protocol Code: MD-20-16254-5).

### Patient recordings and preprocessing

We included microelectrodes recordings from ten patients. All patients had intracranial electrodes placed for presurgical evaluation of epilepsy and the research protocol did not intervene in the placement of electrodes. These participants were included in a project on memory and slow waves in epilepsy, part of which has already been published (Sheybani et al., 2023). Microwire signals were selected based on high signal-to-noise ratio, as reflected by the detection of ≥ 1 single unit. Duration of recordings was of (median, interquartile range) 10 min and 17 s [3-13 min] and number of electrodes per patient was 4.5 [2.75-8].

Behnke-Fried electrodes were inserted in the mesial temporal lobe and signals were recorded at 30 kHz using a Blackrock Neuroport system (Blackrock Neurotech LLC).

Signal artefacts were identified by eye and removed from all analyses. Before FR detection, we notch filtered the data at 50 Hz (49-51 Hz, *bandstop.m* in Matlab). Data were then downsampled to 3000 Hz, 4000 Hz, 5000 Hz, 6000 Hz, 8000 Hz,10000 Hz and 15000 Hz after applying a low-pass anti-aliasing filter at half the new sampling frequency. This was intended to test whether sampling frequency affects the proportion of FRs as distinct phenomena. Prior approval was granted by local Ethics Committee (15/LO/1783).

### FR detection

We used the widely used Delphos detector (Roehri et al., 2016), which has been validated for the detection of FRs (Roehri et al., 2017). The detector measures specific time width and frequency spread of candidate events, which are identified as islands in normalized time-frequency images. This detector has been used in multiple studies focusing on epileptic discharges and high-frequency oscillations (Lambert et al., 2020; Roehri et al., 2018; Schönberger et al., 2021; Shamas et al., 2023). Default settings were used, notably the frequency boundaries for FRs (250-499 Hz). While multiple studies have established that FRs arise from timely generated APs (Foffani et al., 2007; Ibarz et al., 2010; Staba, 2012), ripples reflect post-synaptic potentials (Staba, 2012). This major difference makes the hypothesis that ripples could also emerge from chance firing invalid, explaining why we focus here on FRs generation, not ripples generation.

Detection of FRs in rats could not be visually checked given the large amount of data (c. 11,905 hours of recording). To exclude the possibility that they reflect movement-related artifacts – although Delphos is meant to control for lower frequency artifacts (Roehri et al., 2018) – we computed the proportion of FRs that occur during gamma surges, in both the wake and sleep conditions. To do so, we filtered the signals within the gamma range (20-80 Hz) and estimated dynamic amplitude using the Hilbert transform. We then defined gamma surges as any epoch of 20 ms around EEG amplitude peaks that were greater than two standard deviations above the mean gamma amplitude across the whole recording. A given FR was considered to overlap with a gamma surge if more than 50% of its duration occurred during such epochs. Furthermore, to check the relationship between FR incidence and delta activity, we also identified FRs using a second, fully independent and published HFO-detector (Padmasola et al., 2024; Sheybani et al., 2019, 2018) (see Supplementary Fig. 4c). Briefly, the detector identifies candidate FRs in the filtered data (butterworth, order 2, 250-500 Hz) as oscillations with ≥ 4 cycles whose amplitude is higher than 3 times the surroundings 500 ms baseline. These criteria are based on published literature (Bragin et al., 1999; Lévesque et al., 2011). Candidate FRs that coincided with ripples, high-gamma or gamma activity of a higher intensity are discarded.

To test the circular-to-linear correlation between daily fluctuation of delta amplitude and FR incidence, the phase of delta amplitude fluctuation was extracted for the frequency defined by the peak periodicity, i.e., 24 hours (see Frequency analysis below).

### Simulations

To construct simulations, we first selected one microwire signal based on a lack of movement artefact and presence of > 1 single unit to ensure good signal-to-noise ratio. We then inserted action potentials (APs) corresponding to the mean waveform of all detected APs in Keles et al. (Keles et al., 2024). The number of APs varied depending on the number of simulated neurons (50 steps from 1 to 2000) and mean firing rate (50 steps from 1 to 20) based on realistic firing (Liebe et al., 2025) and including a margin to study what can be expected from higher firing rate. To adjust synchronization, we computed inter-spike intervals (ISIs) using an exponential cumulative distribution function (CDF) and varied the mean parameter μ from 1 – to provide a very exponential distribution of ISIs, i.e., high synchronization – to the total number of APs – to provide an almost linear distribution of ISIs, i.e., low synchronization. Varying synchrony allowed us to test the impact, on the network, of inhibitory cells, which have been shown to favour synchrony (Bocchio et al., 2024; Cobb et al., 1995). APs were then inserted into the signal by convolving the signal with the mean waveform of APs. The amplitude of APs was set to be 5.95 times the standard deviation of background noise, following Keles et al. (Keles et al., 2024).

The computation of simulations follows this step-by-step procedure:

[1] Computation of the mean waveform of all detected APs in Keles et al. (Keles et al., 2024).
[2] Normalization of the mean AP amplitude to 5.95 times the standard deviation of background noise, based on Keles et al. (Keles et al., 2024).
[3] Computation of ISIs

[3a] Computation of the total number of APs as the signal duration * mean firing frequency * number of neurons product
[3b] Computation of ISIs as the exponential CDF, varying the μ parameter, which defines the curvature of the CDF, from 1 (highly exponential) to the total number of APs (almost linear distribution of ISIs).
[3c] Randomization of the ISIs, so that long and short ISIs are not temporally clustered.
[3d] Computation of the number of APs per unit of time, based on the ISIs

This process computes one time series for the entire neural population, thus allowing to precisely control for the inter-neuronal synchronization, a parameter that is crucial in the field of FRs (Ibarz et al., 2010).

The number of neurons recorded by the *in silico* probe was defined by the volume of tissue that can be captured by an electrode’s tip, i.e., 200 μm (Harris et al., 2016). This led to a neural density of 1.73•10^3^ within such a sphere (Henery and Mayhew, 1989; Herculano-Houzel, 2009).

### Gradient boosted tree

To test whether neural count, firing rate and synchronization predict the incidence of FRs, we used a gradient boosted tree (GBT, Matlab function *fitrensemble.m*) with default parameters (method: ‘LSBoost’, number of learning cycles: 100). We compared the actual prediction with the distribution of H_0_, where the number of FRs has been shuffled across neural count, firing rate and synchronization. Across 500 permutations, we saved the coefficient of determination *R^2^* defined by:

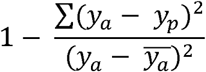

Where *y_a_* is the actual value of FR count, and *y_p_*is the predicted value of FR count. The actual *R^2^* was then compared with the distribution of chance-level *R^2^* for statistics.

To further confirm the performance of GBT, we used a cross-validation procedure where the GBT is trained on 80% of data and then tested on the 20% remaining. The procedure is repeated 1000 times and the *r^2^*is saved at each round. We repeated the analysis with randomization of the outputs across 1000 rounds and saved this null distribution *r^2^*. We then compared the performance against original data.

### Temporal shuffling procedure

To generate time-shuffled EEG signals, we used a wavelet-based iterative amplitude adjusted Fourier transform (wavelet-IAAFT) because it best conserved the spectral content and visual appearance of the original signal (see Fig. 1b for an example and Supplementary Fig. 3 for statistical analyses). Indeed, shuffled signals reliably replicated two key properties of the original signal: mean amplitude (original signal, mean ± SD: 2.5 μV, [±1.2 μV], shuffled signals: 2.7 μV, [±1.3 μV], *p*=0.12, paired t-test) and dominant frequency within the FR range (original signal, median, IQ range: 411 Hz, [399-417 Hz], shuffled signal: 411 Hz, [400-416 Hz], *p*=0.63, Wilcoxon test, Supplementary Fig. 3). Furthermore, this approach allows only preselected frequency components of the original signal to be shuffled, thus controlling for any potential contribution of low-frequency activity to the emergence of FRs (see Fig. 1b, where the low frequency range is conserved, while the high-frequency range is shuffled). Specifically, this reduces the risk of detecting “false-ripples”, i.e., filtered artifacts of lower frequency range (Bénar et al., 2010), although Delphos does control for this kind of artifact (Roehri et al., 2018). This was particularly important for the rat data, where artifacts could not be visually checked (total of c. 11,905 hours of recording).

The algorithm works by decomposing the signal in the frequency domain using a 45-vanishing moment Daubechie wavelet. Then, across coefficients of interest (set at >200 Hz to shuffle only the spectral content that is relevant to FRs), the code extracts the phase of the signal using a fast Fourier transform, generates new (and uniform) phases and then reconstructs the signal iteratively until convergence of amplitude ranks with the original signal. Convergence was assessed using Kullback-Leibler divergence. In animals, shuffling was performed on 1-hour long signals to preserve changes in excitability across time. The code is available on https://github.com/lsheybani/FR_project.

Since the shuffling procedure is central to the present work, we added 5 supplementary ways to randomize time-series. We used Fourier transform (FT)-based phase randomization, amplitude adjusted Fourier transform (AAFT)-based phase randomization, autoregressive-based (AR) randomization of residuals, iterative AAFT until convergence of amplitude distribution (IAAFT-1 in Supplementary Fig. 10) and IAAFT until convergence of spectral power (IAAFT-2 in Supplementary Fig. 10). FT, AAFT, IAAFT-1 and IAAFT-2 are published methods (Lancaster et al., 2018). AR was written in-house. Last, we also implemented a method based on wavelet-IAAFT with preservation of cross-frequency coupling, since fast-ripples are typically locked to low frequency phase (Sheybani et al., 2019). The code detects the highest phase-amplitude coupling (PAC) in the original signal between [300-6000 Hz] for amplitude and several low frequency bands ranging from 2-20 Hz, bandwidth of 3 Hz. PAC is computed using the modulation index (Tort et al., 2008). Then, in the shuffled signal under construction and during convergence testing of PSD (see above), the PAC between high-frequency part of the signal (300-6000 Hz) and the identified low frequency for phase is normalized to that of the highest PAC identified earlier. Parts of this code were written with the help of artificial intelligence (ChatGPT and Copilot).

### Frequency analyses

To compute periodicities of FR incidence and delta amplitude, we first smoothed the data with a sliding window (moving median) of 12 hours to remove small fluctuations in the signals. This was based on visual evidence of slower rhythms for FR incidence and delta amplitude (Fig. 5a). Within animals, we then extracted segments of data for analysis that were at least 48 hours long. We then computed the power spectral density of these signals using Welch’s method (*pwelch* in Matlab) with default parameters (8 segments with 50% overlap, sampling frequency of h^-1^). We proceeded through the same analyses on shuffled data, where incidence of FR or fluctuation of delta amplitude have been randomly permuted across time, 200 times. Periodicities were then identified by plotting the result of the frequency analyses on a 1/frequency axis (i.e., period). We further verified that delta power across time displayed similar fluctuations to beta (15-40 Hz) to delta power ratio, another marker of vigilance (Fraigne et al., 2023).

The time-frequency analysis displayed in Fig. 2c was obtained using a continuous wavelet transform (*cwt.m* in Matlab) and default parameters. For time efficiency, given the large number of simulated EEGs that were generated (125,000) and the consecutively high number of detected FRs, we sub-selected 0.1% of all detected FRs in the simulated signals, which accounted for 5865 events.

### Statistical analyses

We used 2-way ANOVA with Fisher’s LSD post-hoc test and Bonferroni corrections to adjust *p*-values for multiple comparisons. In human subjects, to test for changes in FR rate across sampling frequencies and conditions (original vs shuffled signal), we used a linear mixed model (LMM):

FR incidence ~ 1 + sampling_frequency + condition + sampling_frequency * condition + (1 + sampling_frequency | subjects)

The LMM was performed in Jamovi (version 2.6.22), using the toolbox Gamlj3 (v. 3.4.2). Fixed effects were statistically tested with Omnibus tests.

### Use of Large Language Models

Code for data analyses was written with the help of ChatGPT and Copilot. After using these tools, the authors reviewed and edited the code as needed and take full responsibility for its correctness.

## Declaration of generative AI and AI-assisted technologies in the writing process

In early versions of the manuscript, LS used ChatGPT and Copilot for English editing. After using these tools, the authors reviewed and edited the content as needed and take full responsibility for the content of the published article.

## Data availability

MEA and rodent data can be obtained upon reasonable request from the corresponding author. We will share clinical data on request, provided that request is consistent with the terms of our ethics data collection and analysis.

## Code availability

Code for the project is available from https://github.com/lsheybani/FR_project.

## Acknowledgments

This work was funded by a Guarantors of Brain fellowship and a Swiss National Science Foundation grant to LS (PZ00-3_233452) and supported by the National Institute for Health and Care Research University College London Hospitals Biomedical Research Centre. DB is supported by a UKRI Frontier Research grant (EP/X023060/1).

We are very grateful to Dr. Roman Rodionov for his help in early preprocessing steps of human microwire recordings at UCL. We are very grateful to Dr. Nicolas Roehri for his help in using Delphos for fast-ripples detection.

