## Supplementary Figures for "Fast-ripples are emergent properties of neuronal networks"

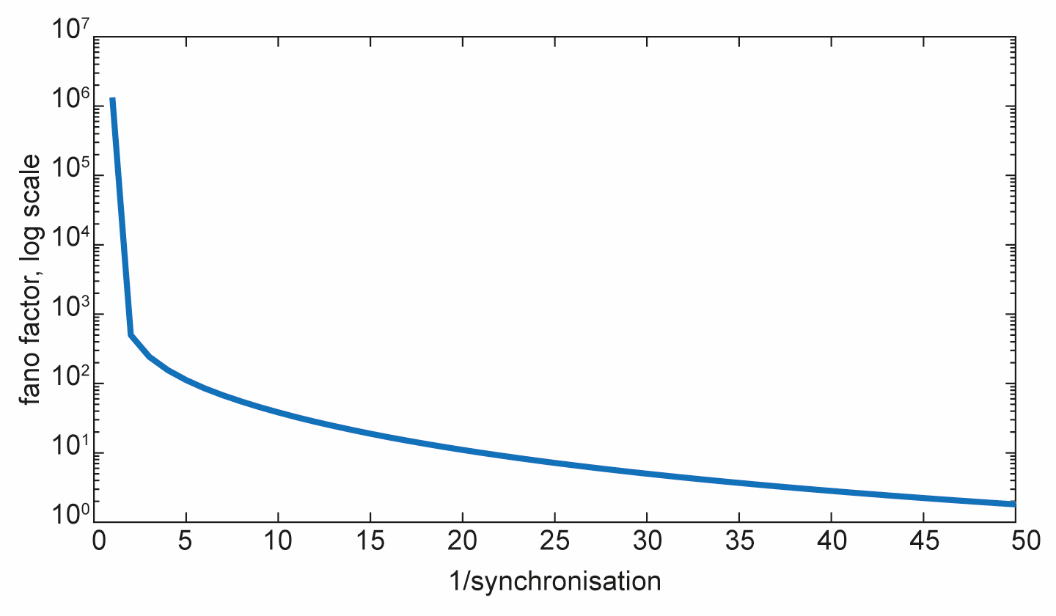

**Supplementary Figure 1 | Fano factor and level of synchronization**

The Fano factor, defined as the variance of interspike intervals divided by their mean, is exponentially related to the level of synchronization of the model.

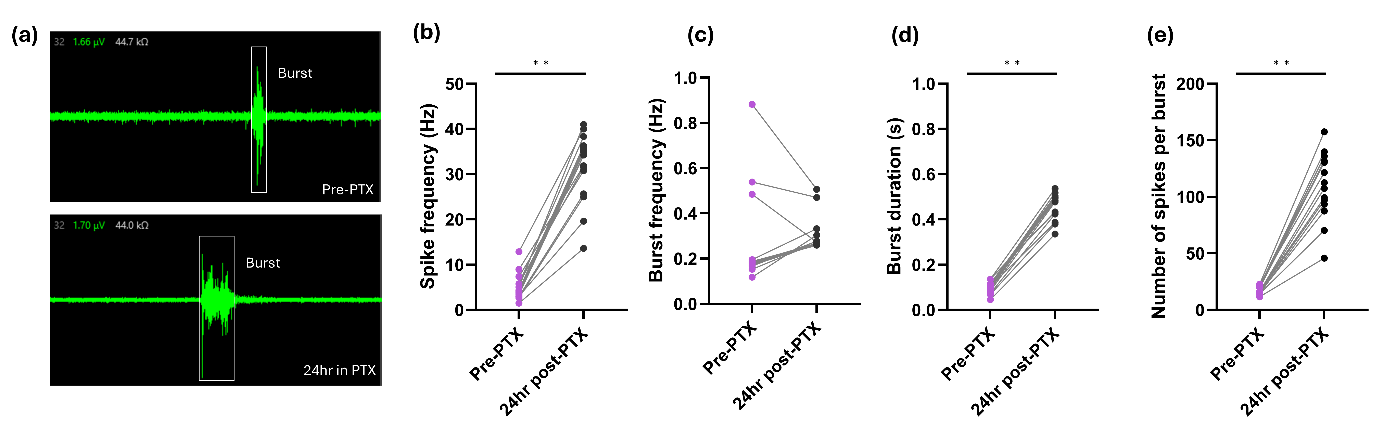

**Supplementary Fig. 2 | Markers of epileptic activity in MEA recordings increase after picrotoxin (PTX) application**

Neuronal subgroup firing events measured by MEA recordings *in vitro*. **(a)** Example waveforms before and after picrotoxin treatment, where a burst event is highlighted. **(b)** Spike frequency, **(c)** burst frequency, **(d)** burst duration, and **(e)** number of spikes per burst (2-way ANOVA, p < 0.001).

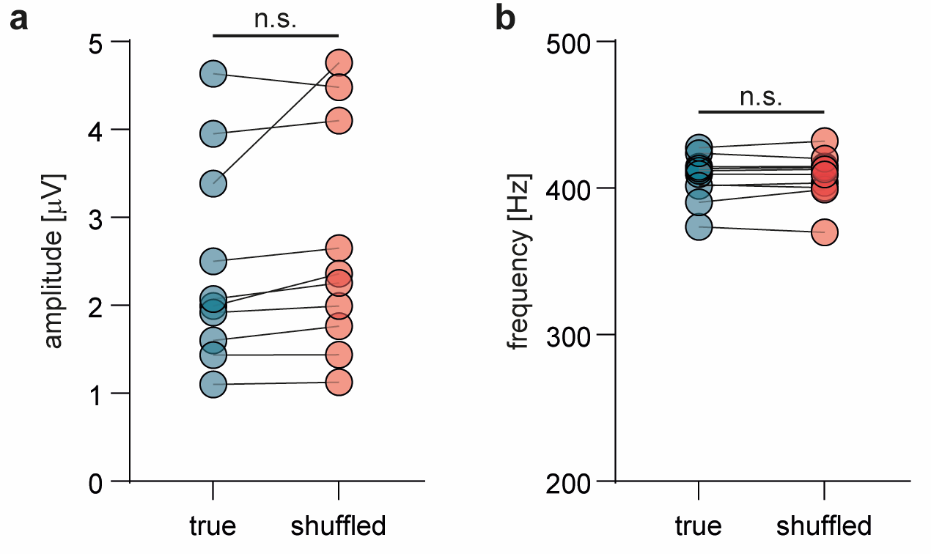

**Supplementary Fig. 3 | Validation of shuffling procedure for high-frequency signals**

To verify that our signal shuffling algorithm did not distort high frequency content, we compared the **(a)** amplitude and **(b)** oscillatory frequency of signals filtered within the frequency range of FRs (200-500 Hz) between the original and shuffled signals. There were no significant differences in either measure. This validation was performed on human data at a sampling frequency of 3000 Hz.

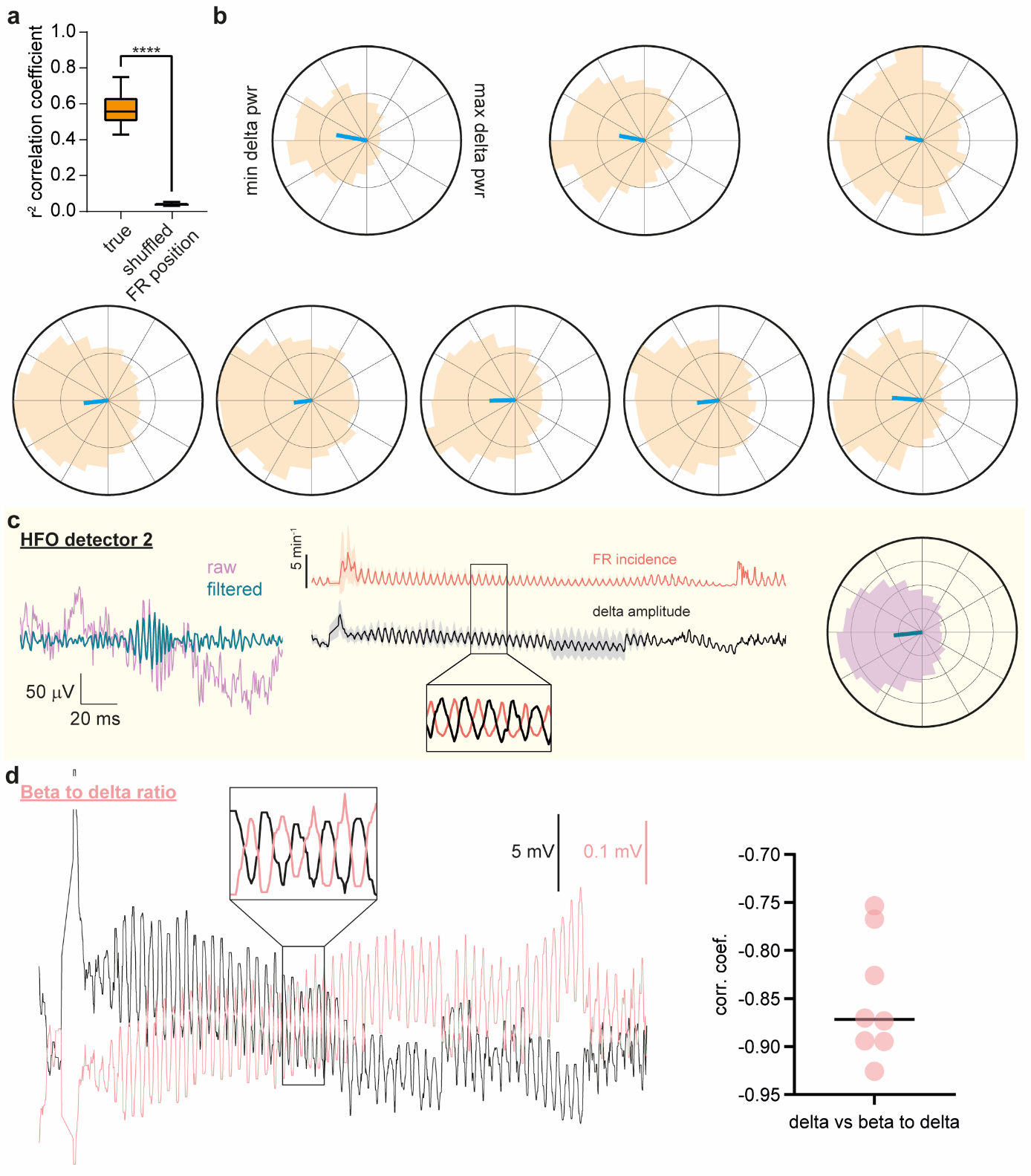

**Supplementary Figure 4 | Distribution of FR incidence along phases of 24-hour delta amplitude fluctuations: group analysis**

**(a)** Circular-to-linear correlation coefficient between the phase of delta amplitude traces and incidence of FRs (left box-and-whiskers). The correlation is significantly higher (r^2^ coefficient, mean, ± SD: 0.57, [0.10]) than if FR occurrence across days is shuffled (r^2^ coefficient: 0.03, [0.01], *p*<0.0001, paired t-test). **(b)** Circular plots indicating the incidence of FRs at different phases of daily delta amplitude fluctuations (-π to π). Each panel corresponds to one animal. The maximal incidence of FRs is seen at the trough of delta fluctuations. **(c)** The inverse relationship between FR incidence and delta fluctuations was further assessed with a second, independent HFO detector. One example FR is depicted (left), as well as the daily fluctuations of FR incidence (middle). Consistent with results from the original detector, FR incidence was inversely proportional to delta amplitude. Hence, FR incidence peaked at delta troughs (right, circular histogram including all animals). **(d)** Beta to delta power ratio across time is superimposed over delta power across time. There is a strong (inverse) correlation between the two time-series (inset), which is confirmed by the correlation coefficient across animals (right).

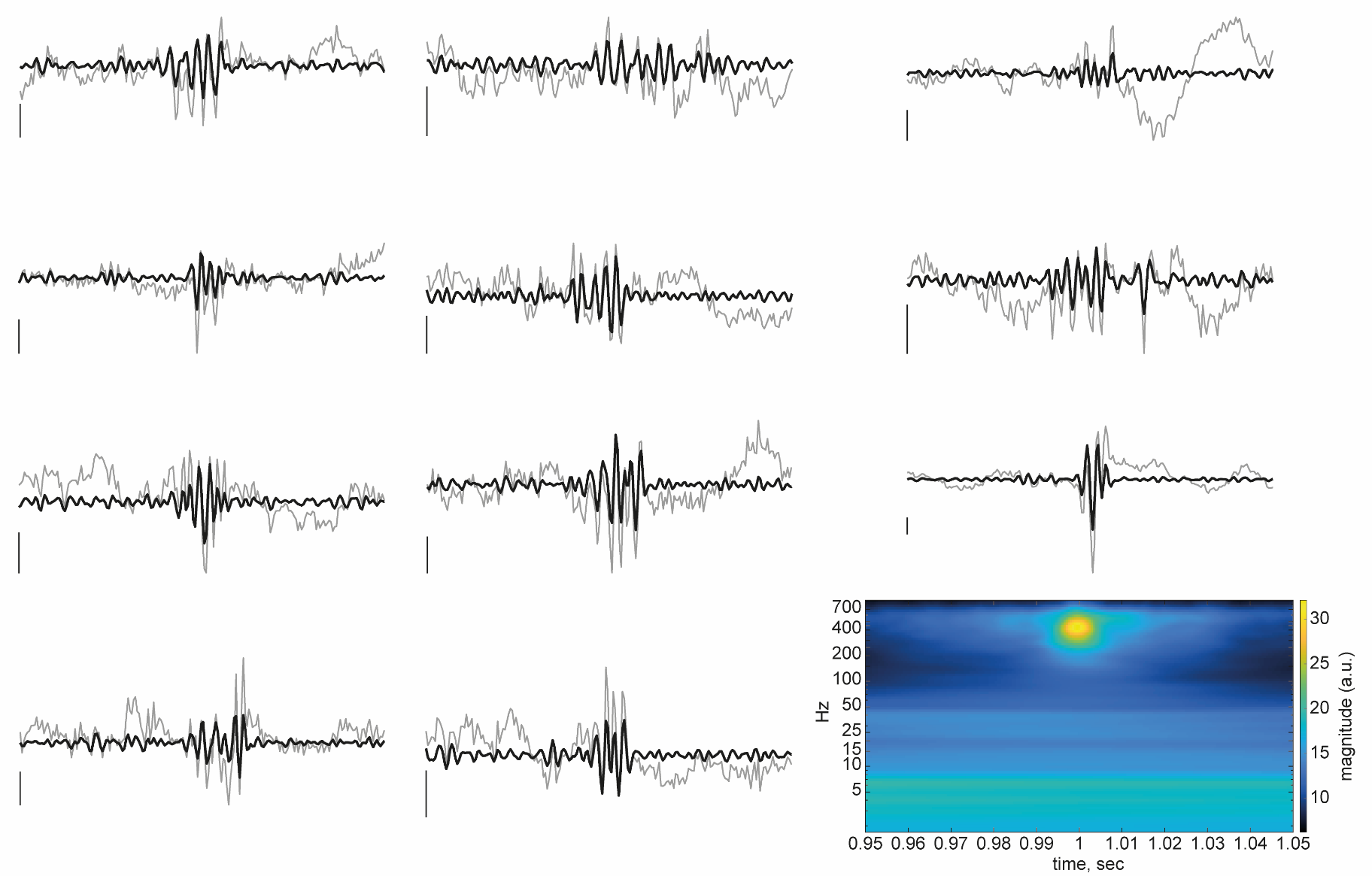

**Supplementary Figure 5 | Examples of FRs in rats**

Examples of FRs in different rats and the time-frequency representation of all detected FRs. See Supplementary Figure 6 for an objective assessment of “false FRs”. Scale: windows of 100 ms, 80 μV.

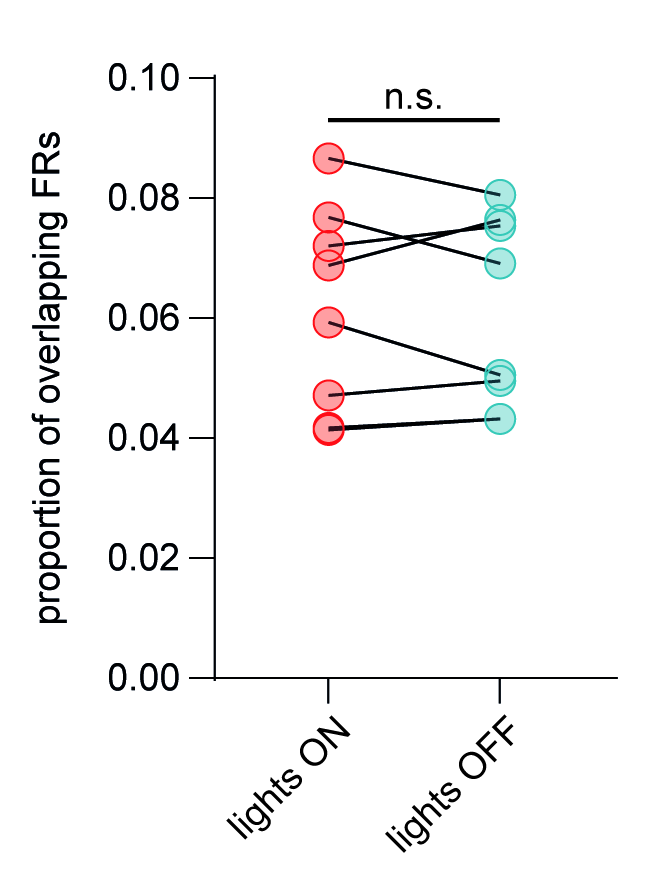

**Supplementary Figure 6 | Lights ON and OFF are not associated with a different proportion of FRs overlapping with muscle artefact (assayed by broadband high-frequency activity)**

We computed the proportion of FRs occurring during periods of broadband high-frequency activity (20-80 Hz) to verify that detected FRs are not “false ripples”. The proportion of such doubtful FRs was very low and, importantly, there was no difference between conditions lights ON and OFF (mean, ± SD, lights ON: 6%, ±2%, lights OFF: 6%, ±2%, *p*=0.74).

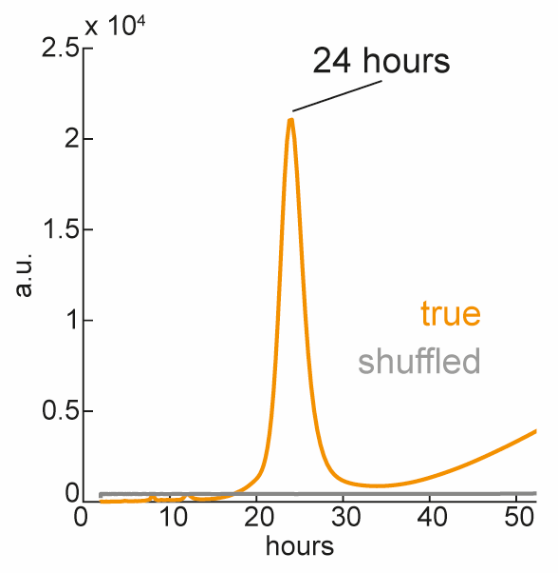

**Supplementary Figure 7 | FR periodogram identifies a peak of incidence each 24 hours**

The periodogram is obtained by computing a frequency analysis over the time-resolved incidence trace of FRs and converting the frequency unit (x-axis) into periods (in hours). A clear peak at 24 hours can be identified, confirming the visually observed daily fluctuation of FRs.

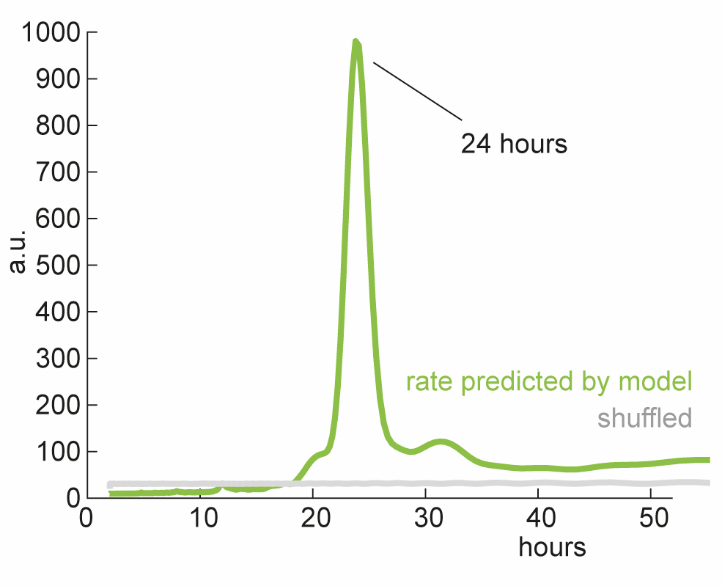

**Supplementary Fig. 8 | Periodogram of FRs incidence in simulated EEGs**

Similar to the rodent data, the incidence rate of FRs in simulated EEGs also displays a period of 24 hours.

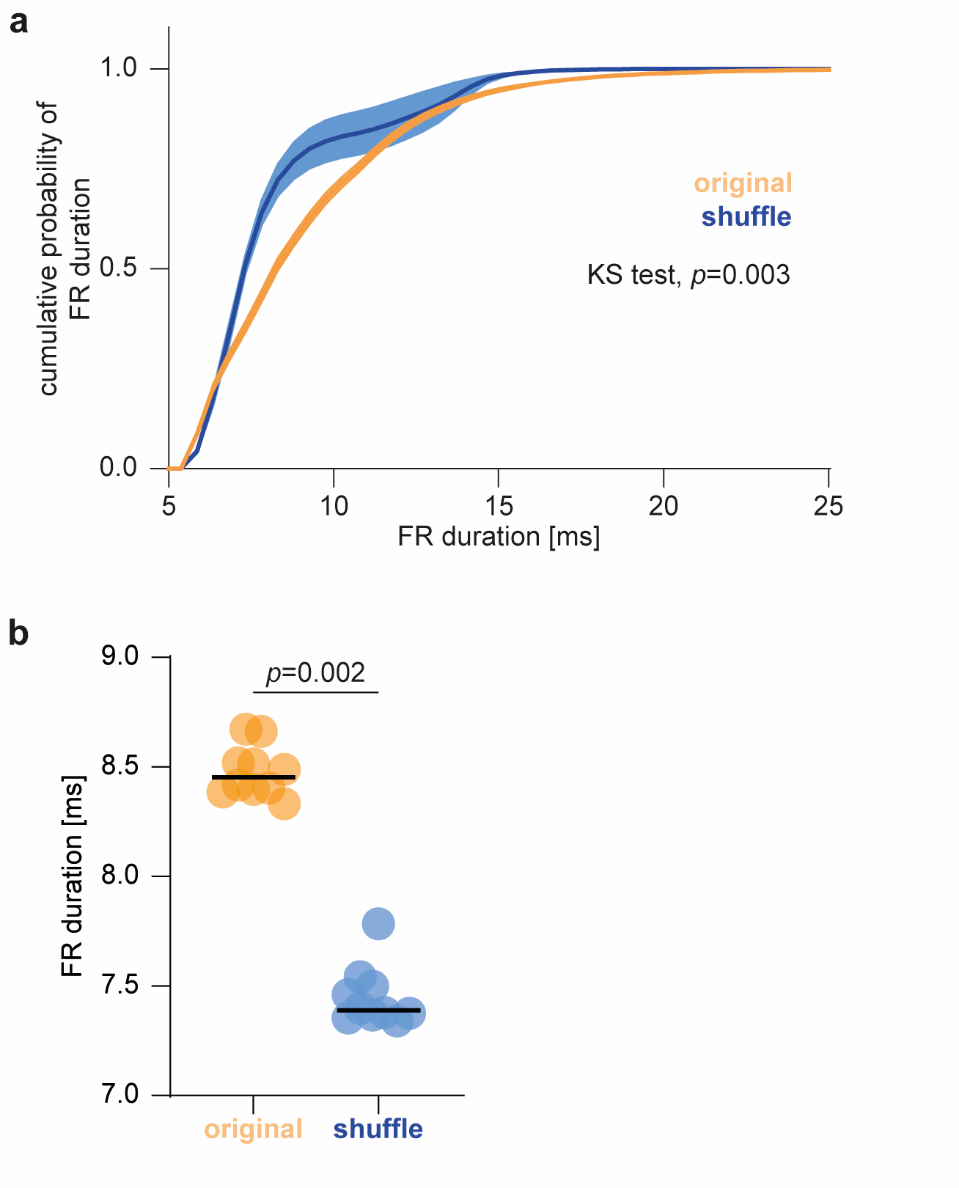

**Supplementary Fig. 9 | Duration of detected FRs**

**(a)** Distribution of detected FR durations in shuffled (blue) and original (orange) rodent data. The two curves differ significantly from 5 to 25 ms (two-sample Kolmogorov-Smirnov test on averages, *p*=0.003). **(b)** The difference observed in (a) is driven by a shorter duration of FRs in shuffled data, as compared to original data in rodents (Wilcoxon test, *p*=0.002).

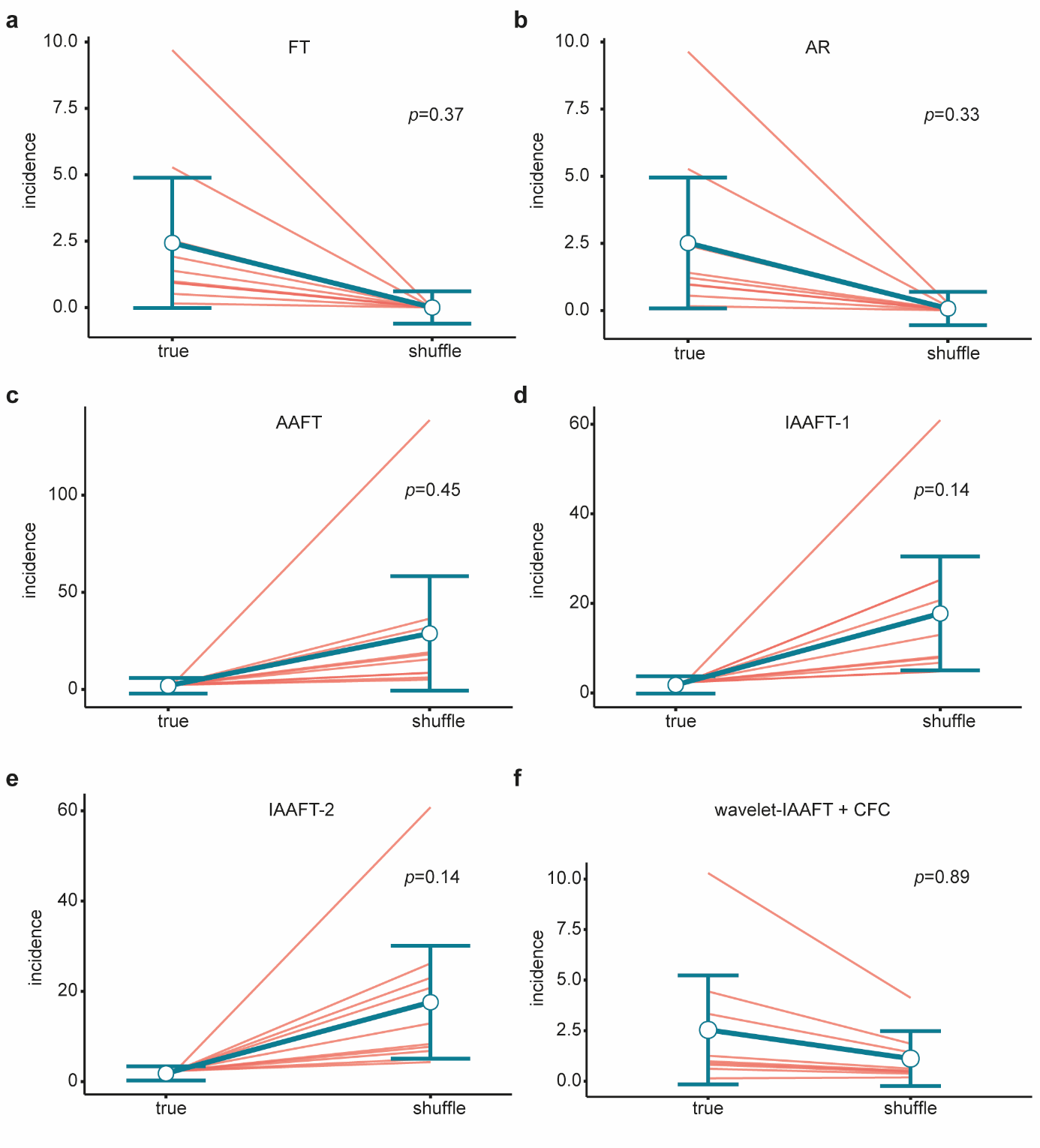

**Supplementary Fig. 10 | FRs are not more frequent than expected by chance when computed with alternative shuffling procedures**

We used a **(a)** Fourier-transform (FT) based, **(b)** auto-regression (AR), **(c)** amplitude adjusted Fourier-transform (AAFT), **(d)** iterative AAFT until convergence of amplitude distribution (IAAFT-1), **(e)** IAAFT until convergence of spectrum (IAAFT-2) and **(f)** wavelet-based IAAFT together with preservation of cross-frequency coupling. None displayed a significantly different incidence of FRs between the original and shuffled data.
